# Low-Cost Tissue Expansion Using a Saline-Based Device Shows Comparable Histological Outcomes in a Rat Model

**DOI:** 10.64898/2026.10.05.756788

**Authors:** Tiago Gonçalves Rosa, Leonardo Carmo Kawakame da Silva, Thiago Zirondi de Abreu, João Stoklos Neto, Jurandir Marcondes Ribas Filho

**Author notes:** Corresponding Author Tiago Gonçalves Rosa, Graduate Program in Principles of Surgery, Evangelical Mackenzie College of Paraná (FEMPAR), Curitiba, PR, Brazil, Fax: N/A, Reprint Requests Tiago Gonçalves Rosa, Graduate Program in Principles of Surgery Evangelical Mackenzie College of Paraná (FEMPAR) Curitiba, PR, Brazil.

## Abstract

Tissue expansion is an established technique for the reconstruction of skin defects, but the cost of conventional silicone expanders may limit its application in contexts with lower resource availability. This study evaluated, in an experimental model, whether a container for storing saline solution could act as a low-cost alternative device for tissue expansion. An experimental, prospective, controlled and randomized study was carried out with 10 Wistar rats, distributed in two groups: conventional silicone expander and saline storage container. Both underwent the same surgical and expansion protocol. Abdominal circumference gain, final expanded skin length, histological and histomorphometric aspects of the skin, including collagen types I and III, as well as exploratory cost analysis, were analyzed. The conventional expander produced a greater gain in abdominal circumference, but the final length of the expanded skin was similar between the groups. Microscopic evaluation showed preservation of the cutaneous architecture, intact epidermis, cellularized dermis, and collagen deposition in both groups. There was no significant difference in the proportion of type I or type III collagen. The saline solution container had a substantially lower cost per centimeter of expanded skin than the conventional expander. These findings suggest that the saline container can produce experimental skin expansion with final length and collagen composition similar to the conventional method, at a lower cost, although additional studies are needed before clinical application.

**Level of Evidence:** Not applicable. Animal experimental study.

## Introduction

The reconstruction of extensive skin defects requires solutions capable of restoring not only the continuity of the skin, but also its barrier function, mechanical resistance, and aesthetic integration with adjacent tissues. Reconstructive surgery has different options within a progressive logic of complexity, from primary closure to grafts, local flaps, and more elaborate tissue transfers. However, these alternatives have predictable limitations, especially in large defects, in which the availability of local tissue is restricted or the morbidity of donor sites becomes clinically relevant (1,2).

Tissue expansion has been consolidated as a unique strategy in this scenario because it allows the gradual generation of autologous tissue adjacent to the defect. Since Neumann’s initial description and the subsequent popularization of the technique by Radovan, cutaneous expansion has come to be understood not only as mechanical distension, but as an active biological process, involving mechanotransduction, cell proliferation, angiogenesis, and reorganization of the extracellular matrix (3-6). Collagen remodeling is a central part of this process, as the ratio between type I and type III collagen reflects important aspects of the maturation, strength, and quality of the expanded tissue (7,8).

However, the widespread adoption of conventional tissue expansion remains limited by economic and operational barriers. Silicone expanders are effective devices, but their cost, the need for prolonged follow-up, and the risk of complications may restrict their use, especially in contexts of lower resource availability (9-12). This reality supports the need to investigate affordable alternatives that preserve the desired biological response, without compromising safety or tissue quality (13).

Based on this gap, this study compared a saline storage container with a conventional silicone expander in an experimental model with Wistar rats. The analysis was structured in three complementary dimensions: macroscopic performance of the expansion, histological and histomorphometric response of the expanded tissue, with emphasis on type I and III collagens, and preliminary economic evaluation related to the cost of the devices (8,14-17).

## Materials and Methods

### Experimental model

This study used a controlled preclinical model, developed to evaluate whether an alternative low-cost device would be able to reproduce the biological and mechanical effects of conventional tissue expansion. Wistar rats were used, a model frequently used in research on wound repair, tissue remodeling, and experimental reconstruction.

### Ethical aspects and study design

The experiment was conducted after approval by the institution’s Ethics Committee on the Use of Animals, under protocol No. 0014/2025. The study followed the institutional standards for the care and use of laboratory animals, as well as the principles of humane treatment applicable to animal experimentation. The study design and reporting were guided, where applicable, by the ARRIVE recommendations for studies involving animals.

The study had an experimental, prospective, controlled and randomized design. Ten Wistar rats were divided into two groups of equal size: conventional expander group and alternative expander group. The same surgical technique, the same postoperative care, and the same expansion protocol were applied to both groups, so that the main experimental variable was the type of device implanted.

### Devices and group allocation

The conventional group received a silicone tissue expander implanted in the subcutaneous plane. The alternative group received a sterile container for the storage of saline solution, used as a low-cost expansion device. The standardization between the groups aimed to allow a direct comparison between the commercial expander and the alternative device. The devices used in the experimental and conventional groups are shown in Figure 1.

**Figure 1.**
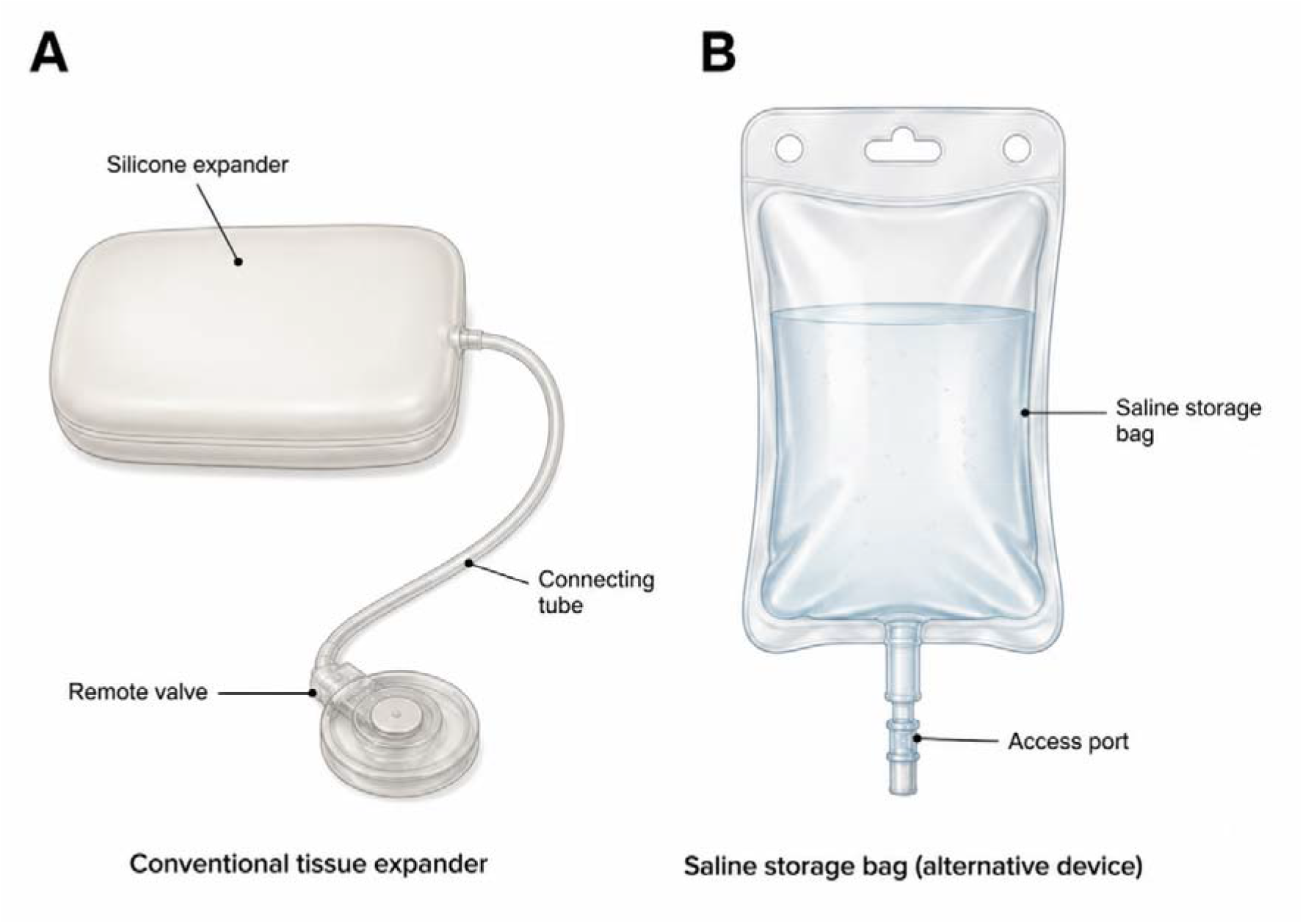
Devices used in the experimental model of tissue expansion. Note: A: Conventional silicone tissue expander with remote valve. B: Container for storing saline solution used as an alternative expansion device.

### Operative technique

All animals were submitted to intraperitoneal anesthesia with ketamine 80 mg/kg and xylazine 10 mg/kg. After anesthetic induction, dorsal trichotomy and antisepsis of the operative area were performed. An incision was made in the dorsal midline, in the interscapular region, followed by blunt divulsion of the subcutaneous plane to create a pocket suitable for the positioning of the device. According to the experimental allocation, the silicone expander or saline container was implanted in the subcutaneous space. The incision was closed with simple nylon suture, and the animals were followed up in the postoperative period for clinical evaluation and detection of possible local complications.

### Expansion Protocol

To allow initial wound healing, tissue expansion was initiated on the fifth postoperative day. The devices were inflated with 10 mL of sterile 0.9% saline solution per day for seven consecutive days, totaling 70 mL per animal. After the end of the expansion, the animals remained under observation for six days for tissue stabilization, and were euthanized on the 18th day of the experiment to collect the biological material.

### Macroscopic and histological evaluation

The macroscopic effect of the expansion was evaluated by measuring the circumference gain and the final length of the expanded skin. After euthanasia, the expanded dorsal skin was resected en bloc and processed for histological analysis. Hematoxylin-eosin staining was used to evaluate the general architecture of the tissue; Gomori’s Trichrome, for analysis of collagen deposition; and Picrosirius Red under polarized light, for differentiation and quantification of type I and III collagens.

### Cost Analysis

A preliminary cost-consequence analysis was performed, comparing the unit cost of the conventional silicone expander with the cost of the saline storage container. The cost per centimeter of expanded skin was also estimated, based on the final length obtained in each group. This analysis was considered exploratory and limited to device cost, and did not represent a complete economic evaluation.

### Statistical analysis

Data were expressed as mean ± standard deviation. Comparisons between the groups were performed by analysis of variance (ANOVA), adopting a significance level of 5%. The absolute gain in abdominal circumference, the percentage gain in circumference, the final length of the expanded skin and the percentages of collagen types I and III were analyzed. The measurements obtained in microscopic fields of the same sample were considered technical repetitions and used to compose the representative value of each animal.

## Results

### Macroscopic outcomes

The two devices showed different behaviors during the tissue expansion process. The conventional expander promoted a greater increase in abdominal circumference, with a mean absolute gain of 8.5 ± 4.7 cm, while the saline solution container resulted in a gain of 4.1 ± 1.3 cm. This difference was statistically significant (p < 0.05). The percentage gain was also higher in Group E, corresponding to 46.4 ± 27.1%, compared to 21.1 ± 7.9% in Group S (p < 0.05). The results of the macroscopic evaluation are presented in Table 1.

**Table 1.**
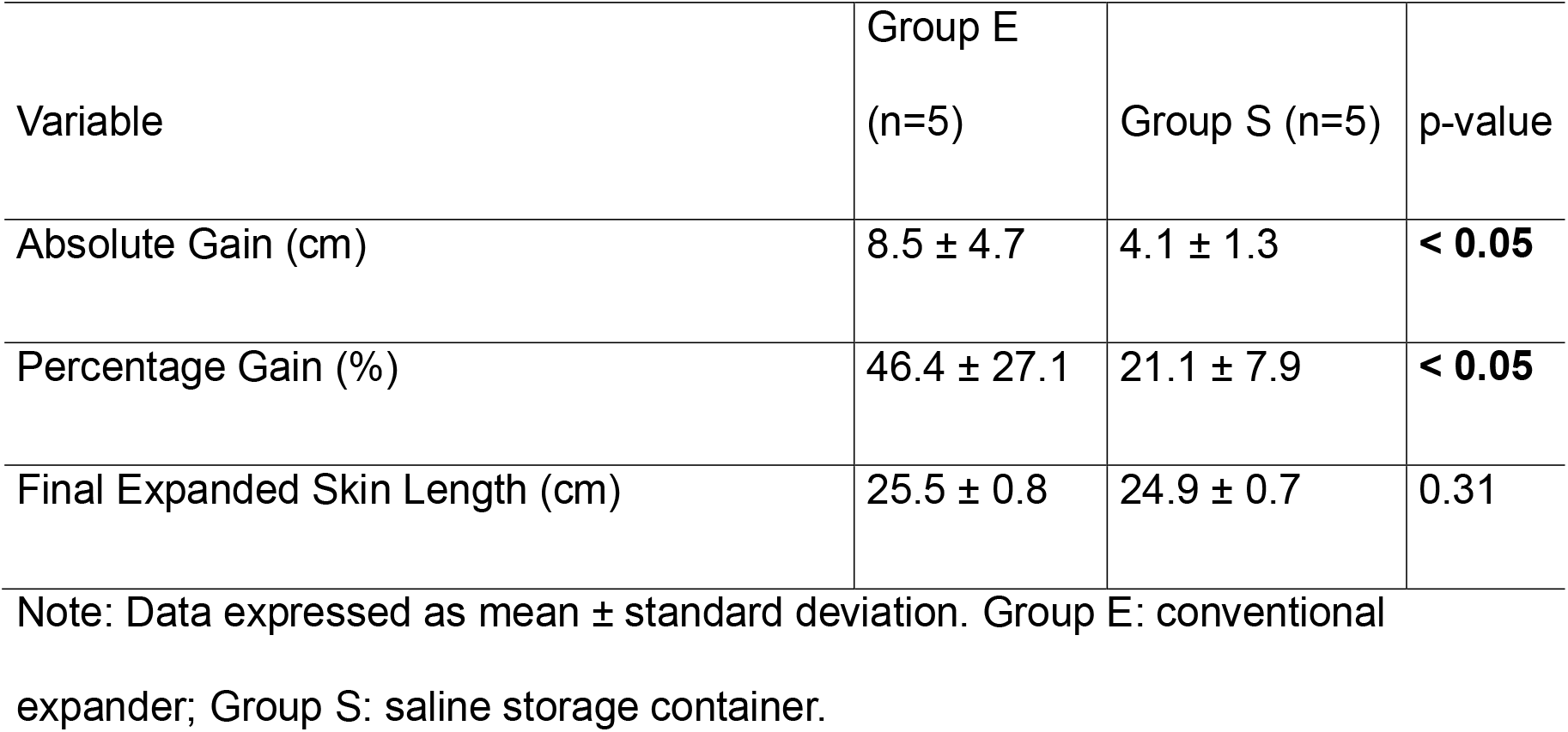
Macroscopic evaluation of tissue expansion in experimental groups.

However, the difference observed in waist circumference was not reflected in a significant difference in the final length of the expanded skin. Group E had a mean length of 25.5 ± 0.8 cm, and Group S, 24.9 ± 0.7 cm, with p = 0.31. Thus, although the conventional expander produced greater circumferential projection, the final amount of skin obtained was similar between the groups.

### Histological and histomorphometric evaluation

Microscopic evaluation demonstrated preservation of the skin architecture in both groups. Hematoxylin-eosin-stained slides showed intact epidermis and dermis with good cellularity. Gomori’s trichrome showed collagen deposition in both groups, while Picrosirius Red under polarized light showed a predominance of type I collagen, with a lower proportion of type III collagen. The histological aspects representative of the expanded skin in the two groups are shown in Figure 2.

**Figure 2.**
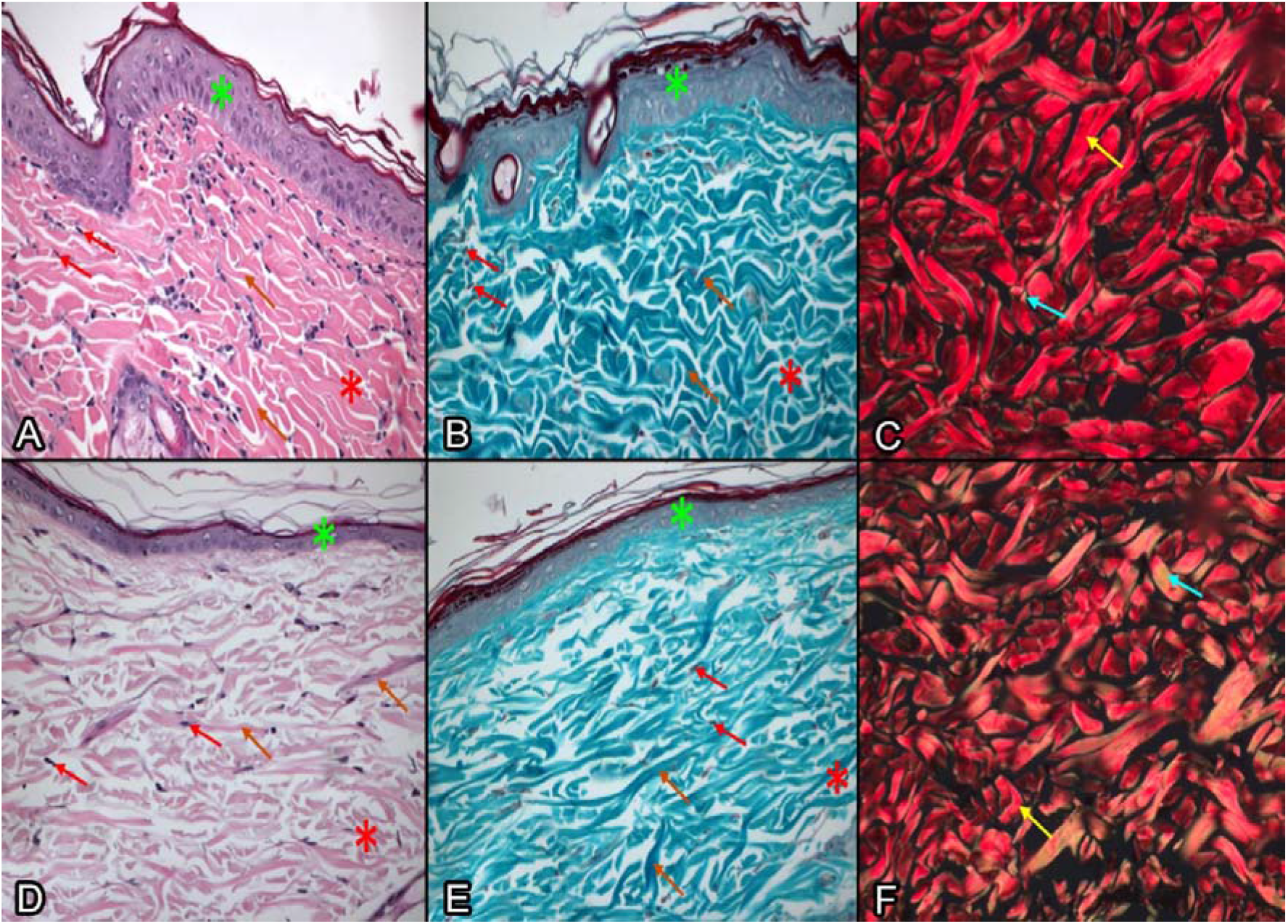
Histological aspects of expanded skin in the conventional and alternative groups. Legend: Histological panel comparing expanded skin. In Group E, a conventional silicone expander was used; in Group S, a saline storage container was used. Hematoxylin-eosin stains demonstrate epithelial preservation and dermal architecture; Gomori’s Trichrome shows collagen deposition; and Picrosirius Red under polarized light shows a predominance of type I collagen, with a lower proportion of type III collagen. Note: HE: hematoxylin and eosin; PSR: Picrosirius Red.

The quantitative analysis reinforced the qualitative findings. The percentage of type I collagen was 93.15 ± 4.24% in Group E and 90.35 ± 5.75% in Group S. For type III collagen, the values were 6.85 ± 4.24% and 9.65 ± 5.75%, respectively. There was no statistically significant difference between the groups for type I or type III collagen, both with p = 0.731. The histomorphometric quantification of type I and III collagen is presented in Table 2.

**Table 2.** Histomorphometric quantification of type I and III collagens in expanded skin.

| Variable | Group E (n=5) | Group S (n=5) | p-value |
| --- | --- | --- | --- |
| Type I collagen (%) | 93.15 ± 4.24 | 90.35 ± 5.75 | 0.731 |
| Type III Collagen (%) | 6.85 ± 4.24 | 9.65 ± 5.75 | 0.731 |

Note: Data expressed as mean ± standard deviation. Quantification performed on slides stained with Picrosirius Red under polarized light. Group E: conventional expander; Group S: saline storage container.

### Economic comparison

The exploratory cost analysis showed a significant difference between the devices. The conventional expander had a unit cost of R$ 5,000.00, while the saline solution container cost R$ 2.50, representing an estimated saving of 99.95% per unit. When the cost was related to the final length of the expanded skin, the value was R$ 196.08/cm in Group E and R$ 0.10/cm in Group S. The exploratory analysis of the cost of the devices is presented in Table 3.

**Table 3.** Exploratory cost analysis of devices per centimeter of expanded skin.

| Parameter | Group E | Group S |
| --- | --- | --- |
| Device Cost (R\$) | 5,000.00 | 2.5 |
| Skin Length (cm) | 25.5 | 24.9 |
| Cost per cm of Skin (R\$/cm) | 196.08 | 0.10 |
| Estimated savings per unit (%) | - | 99.95% |

Note: The analysis considers only the unit cost of the devices and the final length of the expanded skin, and does not represent a complete economic evaluation. Group E: conventional expander; Group S: saline storage container.

The graphical representation of this difference shows the magnitude of the cost reduction associated with the alternative device. The cost per centimeter of expanded skin in each group is illustrated in Figure 3.

**Figure 3.**
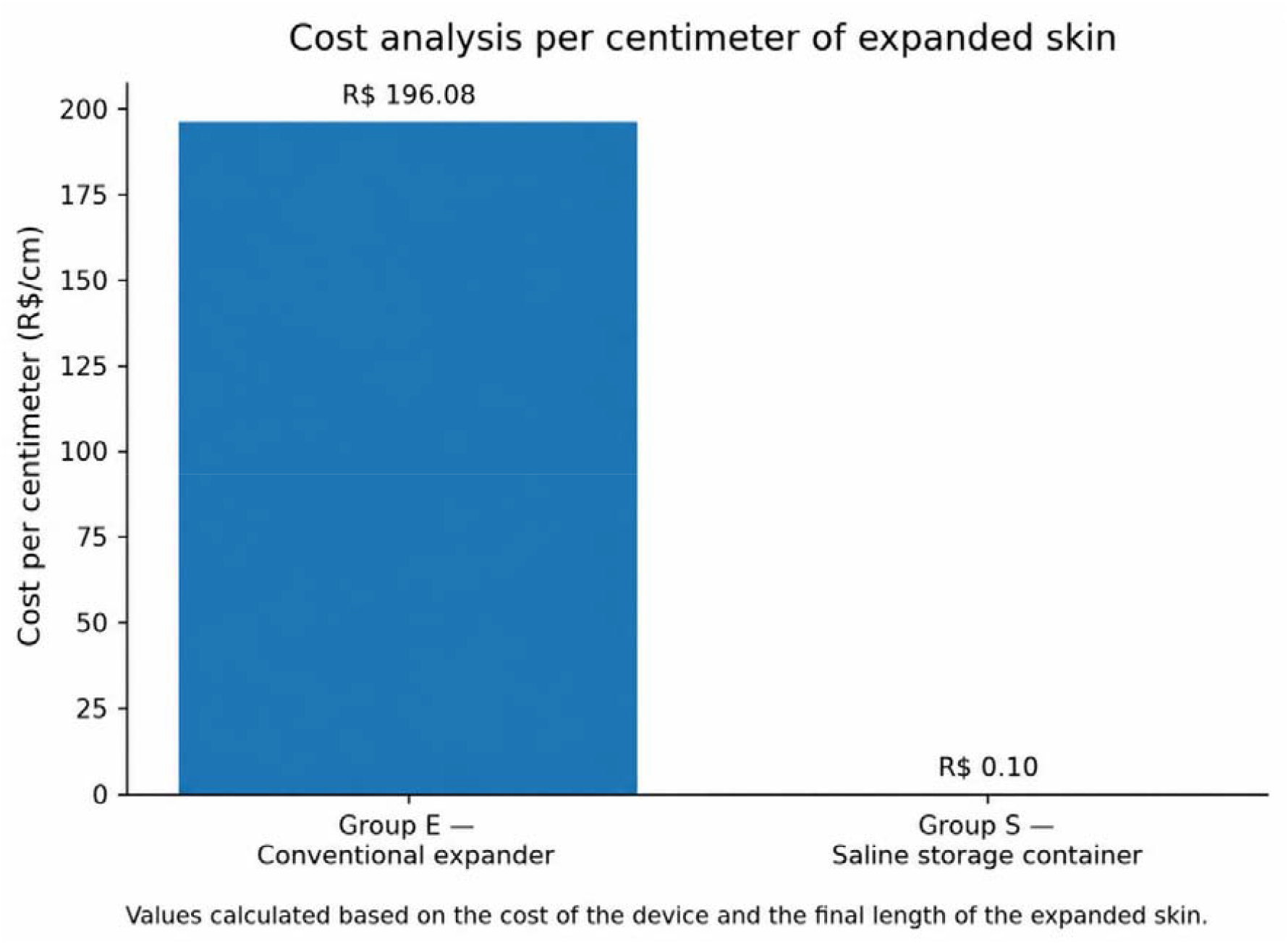
Cost per centimeter of expanded skin in the experimental groups. Legend: Exploratory analysis of the cost per centimeter of expanded skin. The saline solution container presented a substantially lower cost than the conventional expander when adjusted to the final length of the skin obtained.

In an integrated manner, the results indicate that the saline storage container, although it presented a lower circumferential gain, produced a final expanded skin length and histomorphometric pattern of collagen similar to the conventional expander, with a substantial reduction in the cost of the device.

## Discussion

The main findings of this study indicate that a saline solution container can partially reproduce the effects of conventional tissue expansion in an experimental model. Although the silicone expander produced greater gain in abdominal circumference, the final length of the expanded skin was similar between the groups. In addition, the histological and histomorphometric analysis demonstrated a comparable pattern of tissue preservation and deposition of collagen types I and III. These results suggest that the alternative device was able to generate sufficient mechanical stimulus to induce effective skin remodeling.

Tissue expansion is recognized as an active biological process, in which the progressive application of tension triggers cell proliferation, angiogenesis, reorganization of the extracellular matrix, and collagen deposition (5,6,18). In this context, the present study reinforces the hypothesis that the skin’s response to expansion depends less on the sophistication of the device and more on the ability to maintain controlled mechanical tension over time. The similarity in the final length of expanded skin between the groups supports this interpretation.

The greater circumferential gain observed in the conventional expander group probably reflects the greater rigidity and geometric stability of the silicone, characteristics that favor more accentuated local projection. This behavior is consistent with studies that demonstrate that the shape and physical properties of the expander influence the pressure distribution and morphology of the expansion (19). In the group treated with saline solution, the greater malleability of the device may have produced a less projected and more diffuse expansion, without, however, compromising the final amount of skin obtained.

Microscopic analysis adds a relevant element to the interpretation of the results. The presence of preserved epidermis, organized dermis, and absence of exacerbated inflammation indicates that both devices were compatible with the maintenance of tissue viability in the period evaluated. Quantification by Picrosirius Red under polarized light showed a predominance of type I collagen and a lower proportion of type III collagen in both groups, with no statistically significant difference. Since type I collagen is associated with greater maturity and strength of the extracellular matrix, while type III collagen is more related to early phases of repair and remodeling, the similarity between the groups suggests a comparable pattern of tissue maturation (7,8).

These findings dialogue with the literature on cutaneous mechanobiology, according to which mechanical forces are capable of modulating tissue architecture, homeostasis, and reparative response (20). Expanded skin results not only from passive stretching, but from progressive biological adaptation, mediated by fibroblasts, extracellular matrix, and tension-activated signaling pathways (5,18).

Thus, the performance of the saline solution container can be understood as a consequence of a mechanical stimulus sufficient to activate these mechanisms, even if through a device of lower cost and lower rigidity.

Cost analysis amplifies the translational relevance of the study. The markedly lower unit cost of the saline container and the lower cost per centimeter of expanded skin suggest potential for frugal innovation in skin reconstruction. This concept is particularly important in resource-limited health systems, where access to commercial devices may restrict the application of already established reconstructive techniques (12,13). However, this economic advantage should be interpreted as an exploratory analysis, as indirect costs, sterilization, operative time, management of complications, hospital logistics, or regulatory requirements were not incorporated.

The limitations of the study should be considered. The Wistar rat model is suitable for preclinical investigation of healing and tissue expansion, but presents relevant differences in relation to human skin, including thickness, presence of panniculus carnosus, density of appendages, and biomechanical properties (2,15). In addition, the number of animals was small, follow-up was limited to 18 days, and no long-term biomechanical, molecular, or functional evaluations were performed. For these reasons, the results should not be interpreted as a definitive demonstration of clinical equivalence.

Future studies should include larger sample sizes, evaluation of mechanical strength, molecular analysis of the inflammatory and reparative response, prolonged follow-up, and reconstructive models that simulate complex skin defects. Specific investigations on sterilization, material safety, risk of infection, technical reproducibility, and eventual adaptation of the device for clinical use will also be necessary. The translational trajectory should follow principles of progressive evaluation of surgical innovation, avoiding premature extrapolations (21,22).

These findings should be interpreted as early preclinical evidence and as a basis for further studies on alternative tissue expansion devices. Confirmation of the device’s safety, technical reproducibility, expanded tissue stability, and biomechanical performance will be essential prior to any extrapolation to clinical practice.

## Conclusion

The results of this study indicate that the saline storage container was able to reproduce, in an experimental model, relevant aspects of conventional tissue expansion. Despite the lower circumferential gain, the alternative device produced a final length of expanded skin similar to the silicone expander, with preservation of the skin architecture and comparable histomorphometric profile of collagen between the groups. Associated with the significant cost reduction, this finding suggests experimental potential as a low-cost alternative for skin expansion. However, as this is a preclinical study, with a limited sample and short follow-up, additional studies are needed to evaluate safety, biomechanical performance, and clinical applicability.

## Ethical approval

The study was approved by the Institutional Animal Care and Use Committee under protocol No. 0014/2025. All applicable institutional and national guidelines for the care and use of animals were followed.

## Use of artificial intelligence

Generative artificial intelligence tools were used as editorial support for organization, linguistic revision, and adaptation of the manuscript to the journal’s standards. All scientific content, data analysis, interpretation of results, and final approval remained under the full responsibility of the authors.

### List of Abbreviations

AI: artificial intelligence
ARRIVE: Animal Research: Reporting of In Vivo Experiments
CAPES: Coordenação de Aperfeiçoamento de Pessoal de Nível Superior
HE: hematoxylin and eosin
PSR: Picrosirius Red

## Compliance with Ethical Standards

### Informed consent

For this type of study, informed consent is not required.

